# The Toxoplasma Acto-MyoA motor complex is important but not essential for gliding motility and host cell invasion

**DOI:** 10.1101/001800

**Authors:** Saskia Egarter, Nicole Andenmatten, Allison J. Jackson, Jamie A. Whitelaw, Gurman Pall, Jennifer Ann Black, David J. P. Ferguson, Isabelle Tardieux, Alex Mogilner, Markus Meissner

## Abstract

Apicomplexan parasites are thought to actively invade the host cell by gliding motility. This movement is powered by the parasite own actomyosin system and depends on the regulated polymerisation and depolymerisation of actin to generate the force for gliding and host cell penetration. Recent studies demonstrated that *Toxoplasma gondii* can invade the host cell in the absence of several core components of the invasion machinery, such as the motor protein myosin A (MyoA), the microneme proteins MIC2 and AMA1 and actin, indicating the presence of alternative invasion mechanisms. Here the roles of MyoA, MLC1, GAP45 and Act1, core components of the gliding machinery, are re-dissected in detail. Although important roles of these components for gliding motility and host cell invasion are verified, mutant parasites remain invasive and do not show a block of gliding motility, suggesting that other mechanisms must be in place to enable the parasite to move and invade the host cell. A novel, hypothetical model for parasite gliding motility and invasion is presented based on osmotic forces generated in the cytosol of the parasite that are converted into motility.

## Introduction

The phylum Apicomplexa consists of more than 5000 species, the majority of which are obligate intracellular parasites, and includes important human and veterinary pathogens, such as *Plasmodium spp.* or *Toxoplasma gondii* responsible for malaria and toxoplasmosis, respectively. No effective vaccines are available for use in humans and antiparasitic therapies against Apicomplexa are in general of limited efficacy due to drug resistance and lack of potency against chronic stages.

While most microbes rely on the modulation of host cell factors to trigger their uptake via endocytosis or phagocytosis, some apicomplexan parasites, such as *T. gondii* or *Plasmodium falciparum* evolved a highly complex machinery to penetrate their host cell actively within a few seconds. Although alternative mechanisms are described for other Apicomplexans, such as *Theileria spp.* or *Cryptosposridium spp.,* the current theory of active host cell invasion places the actomyosin system of the parasite in the centre of the force that powers gliding motility and host cell entry [1].

The linear motor model of the parasite (Figure 1A) predicts that the myosin A (MyoA) motor, is anchored to the inner membrane complex (IMC) located underneath the plasma membrane and moves on short actin filaments, which are connected via the glycolytic enzyme aldolase to transmembrane and surface-aligned adhesin-like proteins that act as force-transducers [2]. Smooth capping of these force-transducing molecules towards the posterior end of the parasite is associated with forward motion of the parasite on substrates or into the host cell [1]. Additional regulatory components of the MyoA motor complex include the so-called gliding associated proteins GAP40, 45 and 50 and the myosin light chain 1 (MLC1). Based on this model, it is expected that removal of these components would abolish motor function and consequently the parasite ability to locomote or invade the host cell [3–5]. However, video microscopy and biophysical force measurements of gliding *P. berghei* sporozoites suggests a more complex interplay of actin and myosin rather than a linear motor with actin controlling periodic adhesion-deadhesion cycles that account for the stick-and-slip pattern of motility [6; 7].

**Figure 1.**
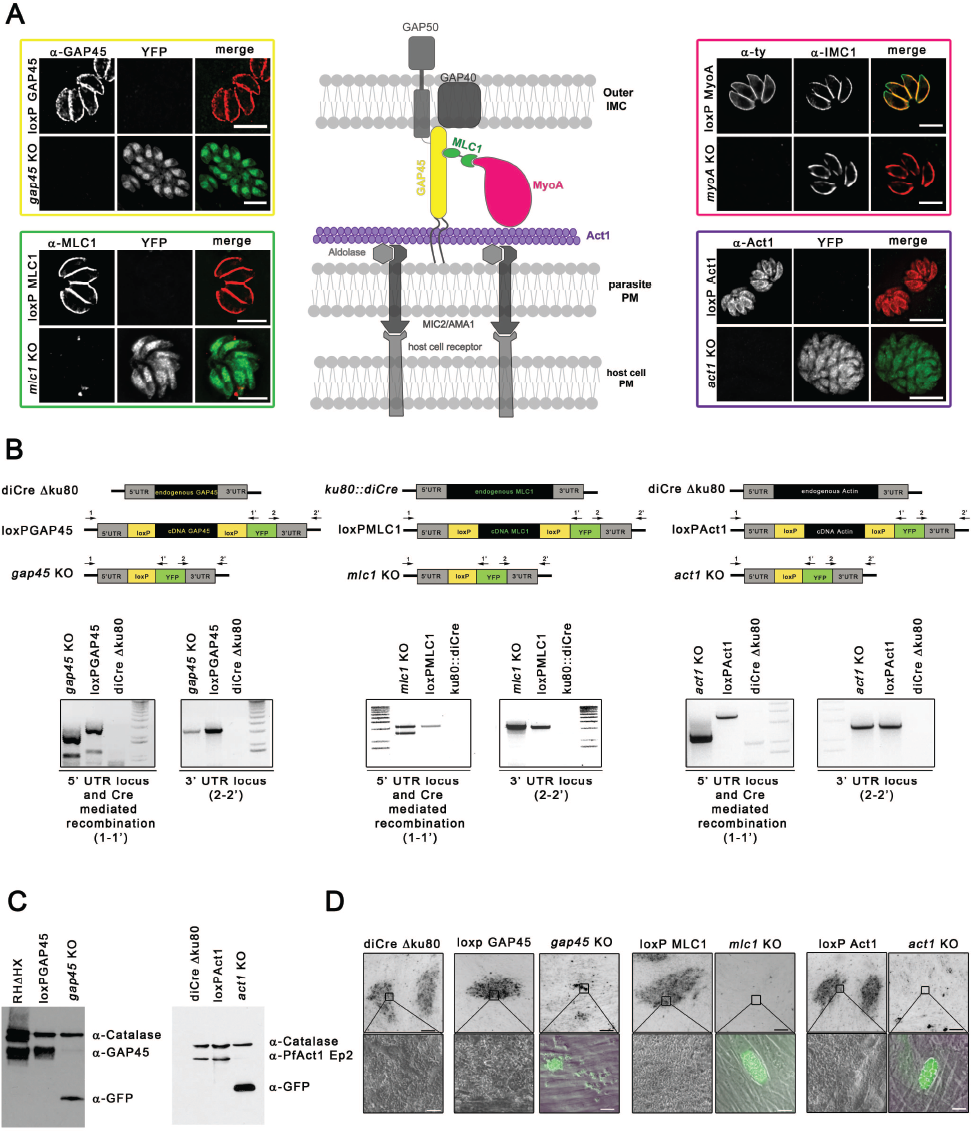
Generation of conditional knockouts for *gap45, mlc1* and *act1*. **(A)** Schematic representation of the apicomplexan gliding and invasion machinery and illustration of the conditional KOs generated for this machinery. Immunofluorescence analysis on the obtained KO strains demonstrate the absence of detectable GAP45, MLC1, MyoA and Act1 respectively using the indicated antibodies. Scale bar: 10 μm. **(B)** PCR on indicated strains shows correct integration of *gap45* cDNA, *mlc1* cDNA and *act1* cDNA into the respective endogenous locus (1-1’; 2-2’). 50 nM rapamycin induction for 4 h shows that Cre mediated excision occurred (1-1’). Primer binding sites are indicated in the schematic **(C)** Immunoblot analysis of GAP45 and Act1 72 hours after excision of the respective gene. Indicated antibodies were used to analyse expression of GAP45, Act1, YFP and as internal control catalase. **(D)** Growth of indicated parasite strains. Plaque formation of indicated parasites was analysed after 5 days. The *gap45* KO, *mlc1* KO and *act1* KO were incapable of plaque formation. Scale bars: 0.2 mm, 20 µm respectively.

Host cell invasion is a multiple step process, consisting of host cell attachment, identification of the host cell by gliding motility, secretion of the rhoptries, establishment of a tight junction (TJ; a ring-like structure at the surface of the host cell) and finally active entry into the host cell through the TJ that serves as a traction point for the actomyosin system of the parasite [8]. Until recently, the traction force potential of the TJ was explained by the formation of a complex between the micronemal protein apical membrane antigen 1 (AMA1) apically secreted and the rhoptry neck protein 2 (RON2), which is a structural component of the TJ inserted into the host cell plasma membrane [9]. Furthermore AMA1 was shown to interact with actin via aldolase fulfilling therefore the requirements for acting as the specific force transmitter during host cell penetration [10]. A detailed reverse genetic analysis of parasites lacking AMA1 has however demonstrated that this protein has no critical role during force transmission, formation of the TJ or parasite entry [11]. Furthermore, preliminary results indicated that parasites lacking MyoA are viable while depletion of Act1 results in loss of the apicoplast but parasites retain some level of invasiveness. These phenotypes suggest that alternative invasion pathways that do not rely on the MyoA-actin motor may operate [12]. These findings suggest several possibilities [12]. First, multiple redundancies can exist that would complement for the removal of individual genes. In case of AMA1 this was considered less likely [11]. Similarly it is hard to imagine a functional redundancy for the single copy gene *TgAct1* [12,13]. In contrast, the huge repertoire of apicomplexan myosins might show some redundancies that could complement for missing MyoA in case of *myoA* KO [12]. Therefore, a more complex invasion mechanism might be in place that can partially substitute for loss of a functional acto-myoA system. In this case one would expect that mutant parasites do not follow the well described step-wise process that includes formation of the typical TJ, a well accepted marker for active parasite entry [14].

To resolve this conundrum we used the DiCre regulation system [12] to engineer parasites lacking proteins of the gliding machinery that are considered as crucial to provide functional motor activity. Conditional knockouts for *gap40, 45*, *50, mlc1*, and *act1* were established and analysed in depth along with the *myoA* KO mutant line previously generated [12]. First we found that parasites without a functional motor complex remained motile indicating that movement can be generated in absence of the known myosin motor and parasite actin. Secondly, none of the generated mutants showed a block in host cell invasion and in all cases entry occurred through a normally appearing TJ. Strikingly, a delay in TJ formation was detected that corresponds to the reduced overall invasion rate of the *act1* KO.

## Results

### Generation of conditional knockouts for mlc1, gap45 and act1

To systematically dissect the role of individual components of the gliding machinery during the asexual life cycle of the parasite, we optimised the DiCre system [12] to generate conditional knockouts (KO) in a novel *DiCre Δku80* strain that shows a significantly higher efficiency of DiCre-mediated recombination upon rapamycin addition (Pieperhoff *et al.,* in preparation). Using this strain, we generated for the first time conditional KO for *gap40*, *gap45, gap50* and *mlc1* as well as a new conditional *act1* KO (Figure 1 and data not shown). As described previously [12], these mutants were generated using the geneswap strategy, where the cDNA of the gene of interest (GOI) is flanked by loxP sides. Upon the Cre-mediated site-specific recombinase activity, the cDNA is removed and the reporter gene YFP placed under control of the endogenous promoter, resulting in green fluorescent KO parasites for the GOI (Figure 1A,B). Efficiency of recombination was monitored based on the percentage of parasites expressing YFP.

All conditional KO were validated on genomic level using analytical PCR with the indicated primer pairs (Figure 1B). Correct 5’ and 3’ integration was verified, and efficient Cre-mediated recombination was achieved in the conditional KO strains. We determined the efficiency of DiCre mediated recombination after addition of 50 nM rapamycin for 4 hours. In the case of *gap45* and *act1* approximately 95% of recombination was obtained (Figure 1B), as only the excised locus was detected in the parasite population using genomic PCRs. In the case of *mlc1*, the recombination rate was lower, with ∼35% of the population losing the gene as judged from analytical PCR showing two bands that correspond to excised and non-excised locus, respectively (Figure 1B).

The absence of the respective gene product in the induced population was also confirmed on protein level in immunofluorescence (Figure 1A) or Western blot analysis (Figure 1C). A faint signal could be observed in the induced populations for GAP45 and Act1 in western blot analysis 72 hours after removal of the respective gene. This corresponds to a minority of parasites that have not excised the floxed gene. Accordingly, in IFA analysis two distinct populations could be identified, with only very few YFP negative parasites (<10%) that were all positive for the protein of interest. In all cases it was straight forward to identify and analyse the respective KO population, since removal of the floxed gene resulted in the activation of YFP expression leading to YFP positive KO parasites. [12] (see Figure 1A,D). Next we performed growth assays on the conditional KO parasite lines, *mlc1* KO, *gap45* KO and *act1* KO Therefore, parasites were induced with 50 nM rapamycin and inoculated on HFF cells. After 5 days the ability to form plaques was analysed and no growth for all three conditional KO lines was observed suggesting that MLC1, GAP45 and Act1 are essential for parasite proliferation *in vitro* (Figure 1D).

Previous attempts to isolate viable KO have been successful for MIC2, MyoA and AMA1 [11,12] while they failed for *mlc1, gap45* and *act1*, confirming the essential nature of the latter genes during the asexual life cycle. We also generated conditional KO for *gap40* and *gap50* (data not shown). In contrast to the other glideosome mutants, deletion of these two genes resulted in an immediate block in intracellular replication of the parasites, demonstrating that individual components of the gliding machinery can have distinct functions during the asexual life cycle. Since this study focuses on the role of the respective genes in egress, gliding and host cell invasion, data on GAP40 and GAP50 will be presented elsewhere.

### Phenotypic analysis of tachyzoites lacking MyoA

It was previously suggested that MyoA is the core motor of the invasion machinery and interacts with MLC1, GAP40, GAP45 and GAP50 [3,4,15]. The phenotypic analysis of the conditional *myoA* KO [12], correlates to the phenotype previously described for a tetracycline-inducible KD mutant of *myoA* where residual host cell invasion was observed but attributed to leaky gene expression of *myoA* [15]. When gliding motility was analysed in a trail deposition assay, *myoA* KO parasites exhibited a residual gliding motility with ∼ 15% of all parasites moving by mostly circular gliding (Figure 2A). Motility of *myoA* KO was further analysed using time lapse microscopy. For this 20 individual parasites were imaged and manually tracked over a time interval of 30 minutes. This kinetic analysis confirmed that *myoA* KO parasites moved slowly for a short distance (Figure 2B,C) and usually stopped after ∼14 µm, for several minutes, before another semicircle was formed confirming the results of the trail deposition assay (supplementary movie 1-2; Figure 2A,B,C).

**Figure 2.**
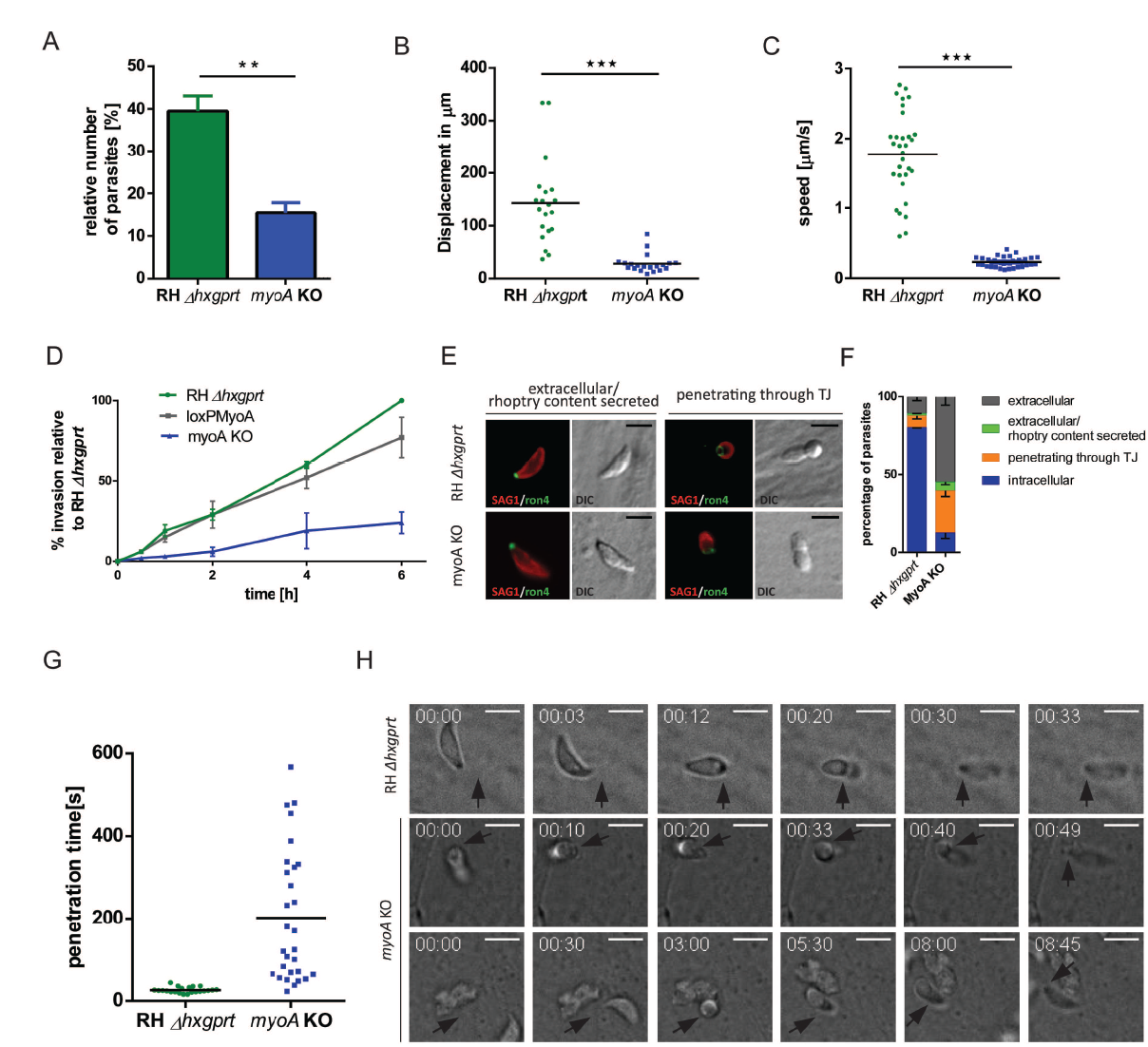
Characterisation of *myoA* KO. **(A)** Quantification of trail deposition assay on indicated parasites. Trails were detected by IFA using antibodies against SAG1. Results are depicted as total % of parasites forming trails. Mean values of three independent experiments are shown, ± SEM **: p-value of 0.007 **(B)** Total displacement of 20 manually tracked parasites for each strain over a time interval of 30 minutes, ***: p-value < 0.001 in a two tailed student’s t-test. **(C)** Graph displays average speed of each motility boost of 20 manually tracked parasites for each strain over a time interval of 30 minutes, ***: p-value < 0.001 in a two tailed student’s t-test. **(D)** Graph shows invasion rates of indicated parasites over time in hours. Data is displayed relative to RH *Δhxgprt* at 6 hours and bars represent ± SEM from three independent experiments. **(E)** IFA against the tight junction protein RON4 and SAG1 in *myoA* KO and RH *Δhxgprt* tachyzoites, scale bar 5 μm. Dot at apical tip of extracellular parasites visualises that rhoptry content has been secreted. **(F)** Graph depicts quantification of extracellular, extracellular and rhoptry content secreted, internalising and intracellular parasites. Data shows mean ± SEM of three independent experiments in triplicate. **(G)** Penetration kinetics of 27 *myoA* KO and 22 RH *Δhxgprt* tachyzoites determined by time-lapse microscopy. Penetration was recorded at one frame per second. **(H)** Time lapse microscopy of RH *Δhxgprt* (upper row) or *myoA* KO (bottom two rows) tachyzoites invading HFFs. Black arrows indicate site of entry, numbers indicate time in minutes:seconds and scale bars represent 5 μm.

Next, we compared invasion and replication rates between *myoA* KO and WT parasites. First, parasites were allowed to invade human foreskin fibroblasts (HFF) for different times and let to replicate for 24 hours. While no significant difference in replication was found, at each time point, invasion rate by *myoA* KO parasites remained constant at ∼25% compared to wt parasites (Figure 2D), indicating that *myoA* KO parasites require more time to reach a similar number of invasion events as compared to RH*Δhxgprt*. For example the invasion rate of *myoA* KO parasites after 2 hours corresponded to the invasion rate of RH *Δhxgprt* after 30 min (Figure 2D). Similar effects were observed using other host cells, such as HeLa cells (not shown). The overall reduction in invasion could be due to several effects. Since one of the earliest markers of the entry process is the TJ [16,17], it was important to assess if the observed reduction in invasion rate is caused by a delay in TJ formation or by a block in parasite progression into the host cell after TJ formation. While *myoA* KO parasites invaded the cell via a normally RON-shaped TJ (Figure 2E), it was found in a 5 min synchronised invasion assay [18], that the majority of *myoA* KO parasites (∼60%) remained attached to the host cell without forming a TJ. Approximately 30% of *myoA* KO parasites formed a TJ and another 10% of all parasites were internalized. In contrast, the majority of control parasites (80%) were found to be intracellular, 10% in the process of entry and only 10% still extracellular without initiation of TJ formation (Figure 2F). Together these data demonstrate that a step upstream of TJ formation, such as host cell recognition or reorientation of the parasites is delayed.

To directly analyse the effect of MyoA depletion on host cell entry, the kinetics of this invasion step was analysed using time-lapse microscopy. In total 22 entry events for control and 27 for *myoA* KO parasites were compared (Figure 2G). As expected, control parasites moved in within 20-30 seconds in a smooth and uniform movement (supplementary movie 3). In contrast, *myoA* KO parasites showed a huge variability, with some parasites penetrating the host cell rapidly in a smooth process. However, the majority entered in a spasmodic stop-and-go fashion, and appeared stalled for several seconds to minutes (see supplementary movies 4-5; Figure 2G, H). The fastest recorded entry was ∼25 seconds, whereas the slowest entry took almost 10 minutes until the parasite was completely internalised (Figure 2H). While these results demonstrate an important function of MyoA for efficient, smooth host-cell penetration, the fact that *myoA* KO parasites remain capable of penetrating at similar speed as wild-type parasites indicates that the force for host-cell penetration can be generated independently of MyoA.

Together these results were interpreted that MyoA plays an important but not essential function in multiple steps during host-cell invasion. Since deletion of other components of the invasion machinery (MLC1, GAP45 and Act1) impeded mutant survival (Figure 1), we speculated that the function of MyoA might be partially complemented by other myosins.

### Overlapping functions of MyoA and MyoB/C

Based on its close homology to MyoA [19], we speculated that MyoC might be capable of complementing for MyoA. Therefore, a conditional triple KO for *myoA* and *myoB/C* (note that MyoB and C are isoforms generated by alternative splicing [20] was generated by removing *myoB/C* in loxPMyoA (Figure 3A). The expected genetic modifications were verified using analytical PCR on genomic DNA (Figure 3A). No growth effect was detected for loxPMyoA-*myoB/*C KO parasites, indicating that *myoB/C* is not important for parasite survival *in vitro* (Figure 3B). Induction of DiCre with 50 nM rapamycin allowed efficient removal of *myoA* (∼60% of the population, data not shown), resulting in *myoA/B/C* KO parasites. Several attempts to isolate a viable triple knockout failed, indicating that removal of all three myosins is not tolerated by the parasite. Indeed, in growth assays it was observed that parasites did not form plaques in a HFF monolayer (Figure 3B). It was found that host-cell egress is completely blocked (Figure 3C). In contrast, albeit ∼5-fold reduced when compared to *myoA* KO (Figure 3D), parasites without *myoA,B,C* remained capable of invading the host cell through a normal appearing TJ (Figure 3E) indicating that MyoA and MyoB/C have overlapping functions.

**Figure 3.**
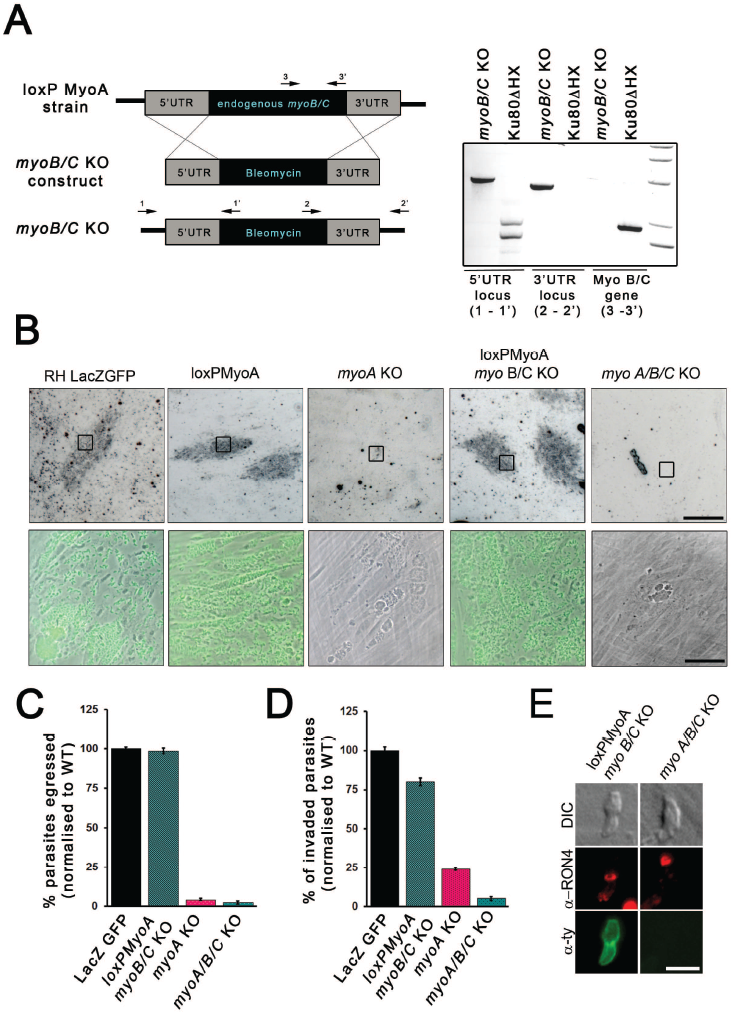
Generation and characterisation of a *myoA/B/C* KO. (**A**) Schematic of loxPMyoA *myoB/C* KO strain. The *myoB/C* gene was replaced for a Bleomycin selection cassette in loxPMyoA strain by homologous recombination. Analytical PCR with indicated oligonucleotides on genomic DNA confirmed the correct integration of Bleomycin (1-1’;2-2’) and absence of *myoB/C* was demonstrated (3–3’). (**B**) Growth analysis of indicated parasites on HFF monolayer for 5 days. To obtain *myoA/B/C* KO, loxPMyoA *myoB/C* KO was treated with 50 nM rapamycin for 4 h prior to inoculation. RH LacZ GFP and loxPMyoA served as controls and showed normal growth behaviour. No growth defect was observed for loxPMyoA *myoB/C* KO. In contrast *myoA/B/C* KO showed a more severe effect on plaque formation compared to *myoA* KO. Scale bars: 0.2 mm; 20 µm respectively. (**C**) Egress analysis of indicated parasites treated ± 50 nM rapamycin 96 h prior to artificial induction of egress with Ca^2+^ ionophore (A23187) for 5 min. For quantification mean values of three independent experiments are shown, ± SEM. The egress rates were standardised to RH LacZ GFP. (**D**) Invasion assay of indicated parasites treated ± 50 nM rapamycin 96 h prior to invasion. Parasites were allowed to invade for 30 minutes before fixation. For quantification 300 parasites were analysed. Mean values of three experiments are shown, ± SEM. All strains were standardised to RH LacZ GFP. (**E**) Immunofluorescence analysis with indicated antibodies on invading parasites demonstrates that parasites without *myoA,B,C* KO form a normal appearing tight junction (TJ). Scale bar: 5 µm.

However, since it is technically not feasible to generate multiple KO for the entire repertoire of myosins that can potentially complement for MyoA, MyoB/C, we characterised other core components of the gliding and invasion machinery.

### MLC1 is essential for host cell egress but not invasion

To date MLC1 is the only of the 7 myosin light chains (MLC) identified to interact with MyoA [21] and GAP40, GAP45, GAP50 in the gliding machinery and is a target for development of invasion inhibitors [5,22]. It was impossible to isolate viable *mlc1* KO parasites, indicating an essential function of this gene. While MLC1 was not detectable in YFP-expressing parasites as soon as 48 hours post induction, the phenotypic analysis depicted in Figure 4 has been performed 96 h after DiCre induction to ensure absence of MLC1. Furthermore, it was possible to maintain *mlc1* KO parasites for up to 2 weeks in the induced population when artificially released from the host cell, before they were overgrown by parasites where no gene excision occurred (not shown).

**Figure 4.**
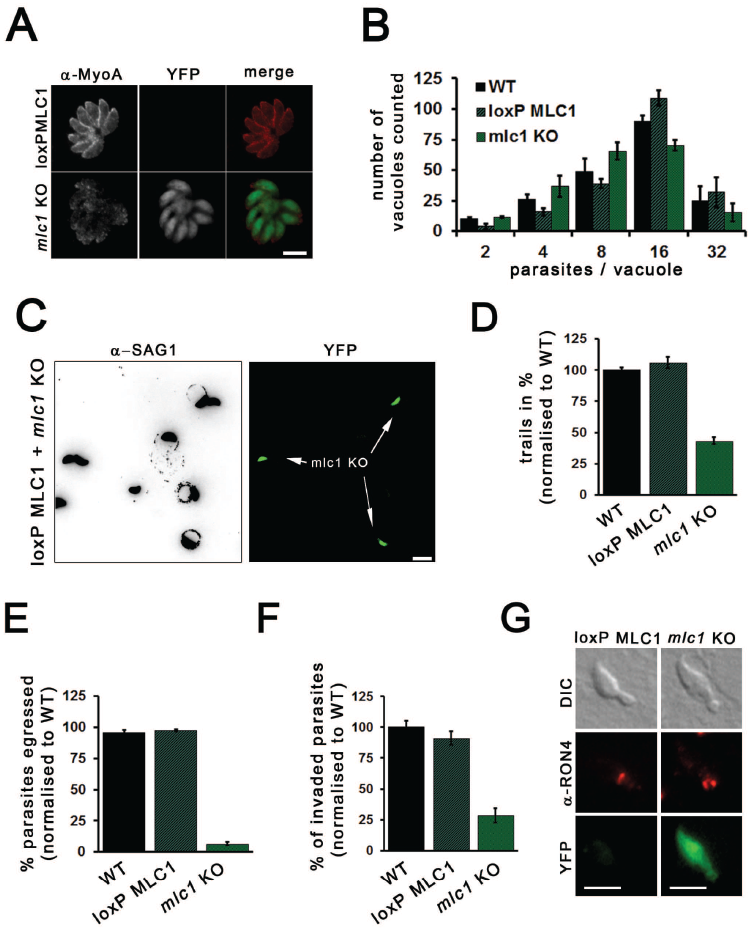
Characterisation of a *mlc1* KO. (**A**) The localisation of MyoA in *mlc1* KO was analysed using α-MyoA (red) antibody 48 hours after induction of loxP MLC1. Loss of MLC1 results in weak punctuated stain for MyoA mainly in the cytosol of the parasite. Scale bars represent 5 µm. (**B**) Replication analysis of *mlc1* KO parasites was performed 96 hours post induction. Parasites were given 1 h to invade prior replication for 24 h. To quantify the replication rate the number of parasites per parasitophorous vacuole was determined. Mean values of three independent assays are shown ± SEM (**C**) Trail deposition assays with indicated parasites were performed by allowing the parasites to glide on FBS coated cover slips for 30 minutes prior to staining with α-SAG1 antibody. The assays were performed 96 hours post induction. Despite the low excision rate of loxP MLC1, *mlc1* KO parasites can be identified easily due to their YFP expression (see arrows). Scale bar: 10 µm. (**D**) Quantification of trail-deposition assays on indicated parasites demonstrates that gliding is significantly reduced in *mlc*1 KO parasites (42% compared to wild type parasites). Only YFP positive parasites were considered to analyse *mlc1* KO. Mean values of three independent experiments are shown, ± SEM. The number of trails was standardised to wild type parasites. (**E**) Egress of loxPMLC1 and *mlc1* KO parasites (treated with 50 nM rapamycin 96 h prior performing egress assays) after artificial induction of egress with Ca^2+^ ionophore (A23187) for 5 min. For quantification of parasite egress mean values of three independent experiments are shown, ± SEM. The egress rates were standardised to wild type parasites. (**F**) Invasion assays of loxPMLC1 and *mlc1* KO parasites were performed 96 h after treatment with ± 50 nM rapamycin. Parasites were allowed to invade for 30 minutes before fixation. Mean values of three independent assays are shown ± SEM. (**G**) Immunofluorescence analysis of invading parasites demonstrates that parasites lacking *mlc1* were capable of invading the host cell through a typical tight junction (TJ). α-RON4 antibody was used to stain for the TJ. Scale bar: 5 µm.

We found that depletion of MLC1 had a direct effect on the localisation and expression level of MyoA as assessed by fluorescence in situ signals (Figure 4A). Only a dotted, weak signal of MyoA could be observed within the cytoplasm of the *mlc1* KO intracellular parasites whereas MyoA delineated the periphery of MLC1-positive parasites (Figure 4A). This cytosolic signal appears to be a cross reaction, since it can be also detected in *myoA* KO (Figure S1) raising the possibility that MyoA is completely degraded in absence of MLC1, as observed in case of *P.berghei,* where depletion of MTIP (the homologue of MLC1) results in complete loss of MyoA [23]. We found no effect during replication after deletion of *mlc1* (Figure 4B). Next we investigated if loss of MLC1 has an impact on gliding motility. For this we only scored trails clearly deposited by YFP-expressing parasites, where *mlc1* has been excised (Figure 4C). Interestingly, mainly circular trails were observed, similar to the phenotype of *myoA* KO parasites. Gliding rate of *mlc1* KO parasites was approximately 42% when compared to controls (Figure 4D). Interestingly, while parasites without MLC1 were unable to egress from the host cell when artificially induced with Ca^2+^-Ionophore (Figure 4E), invasion rate of *mlc1* KO was 28% that of controls (Figure 4F). As expected, *mlc1* KO parasites invaded through a normally appearing TJ (Figure 4G).

In summary *mlc1* is essential for host cell egress, localisation of MyoA at the IMC, but not gliding motility and invasion. Together these data suggest that at least for MyoA function and localisation no other MLC can complement. However, it cannot be excluded that a different motor complex, such as the recently described MyoD-MLC2-motor [21] can partially complement in absence of the MyoA-motor.

### GAP45 is a structural component of the IMC that is dispensable for gliding motility

GAP45 has been described as the membrane receptor for the gliding machinery [4] and its knockdown results in the relocalisation of the gliding machinery to the cytosol of the parasite in conjunction with the loss of IMC stability [3], Surprisingly some parasites remain invasive and reach relative invasion rates of 25% [3].

Since residual expression of GAP45 could not be ruled out in the KD strain, we generated an almost pure parasite population lacking GAP45 following DiCre-induced gene excision (Figure 1A, C) and were able to keep it in culture for up to 14 days. However we did not succeed to isolate a viable *gap45* KO clone. We confirmed that removal of GAP45 resulted in cytosolic localisation of other components of the gliding machinery, such as MLC1 or MyoA [3] (Figure 5A). When compared to depletion of MyoA/B/C or MLC1, depletion of GAP45 impacted tachyzoite morphology. After exiting from their host cells, tachyzoites shifted from a crescent to a round shape, a change that was associated with the IMC detachment from the plasma membrane as observed in ultrastructural analysis (Figure 5B). This crippled overall morphology recapitulated what has been described for *gap45* KD [3] to a more pronounced extent. In contrast, intracellular parasites developed normally and showed no reduction in replication rate, as described before for *gap45* KD parasites [3] (Figure 5C).

**Figure 5.**
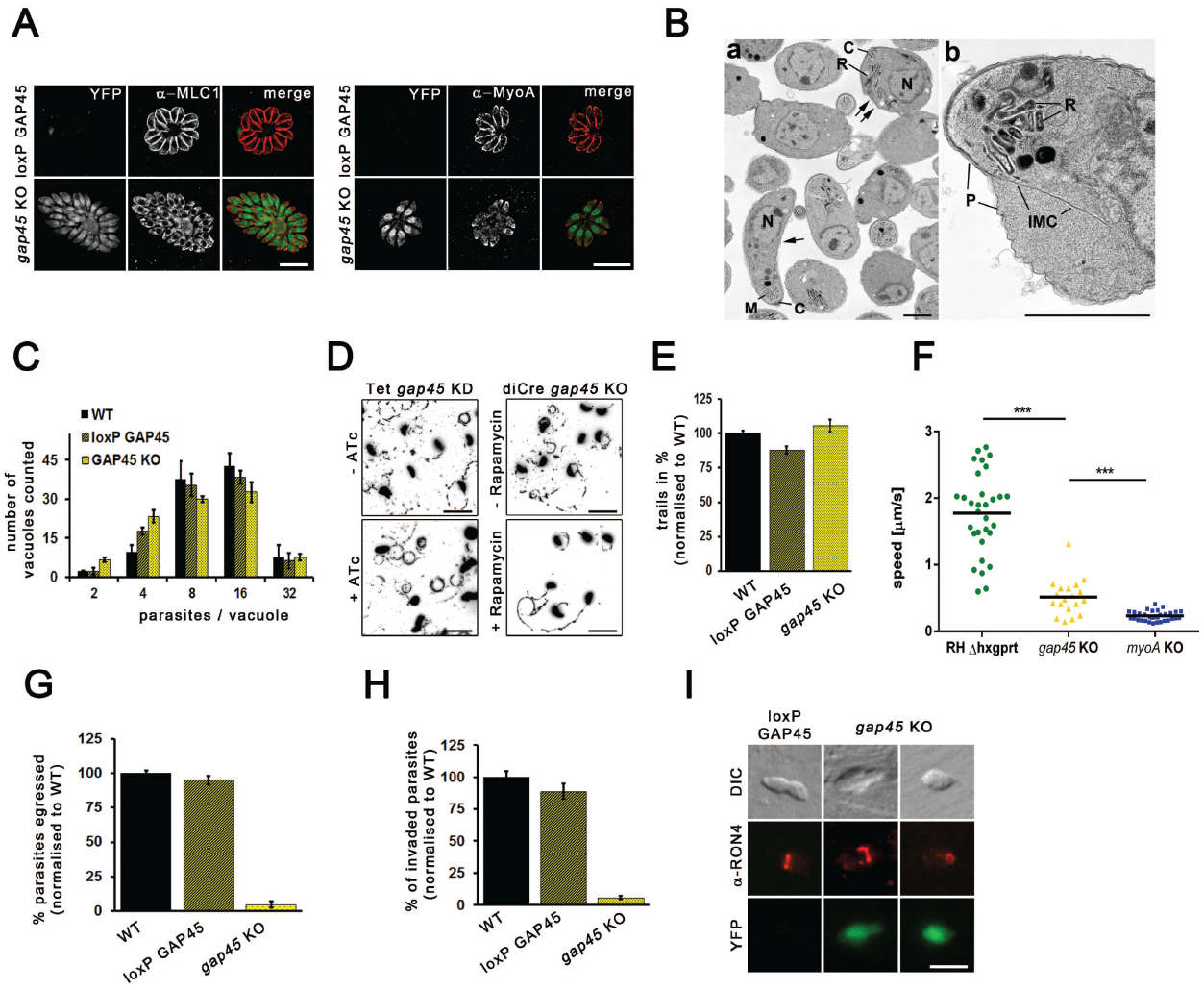
Characterisation of a *gap45* KO. (**A**) Immunofluorescence analysis of *gap45* KO using indicated antibodies 72 hours after induction of loxP GAP45. Both MyoA and MLC1 are re-localised from the periphery of the parasite to the cytosol. Scale bars represent 5 µm. (**B**) Ultrastructural analysis of extracellular *gap45* KO parasites. (a) Low power of *gap45* KO extra-cellular tachyzoites showing the marked variation in shape from normal crescentic shaped parasites (arrow) to more spherical shape parasites (double arrows). The conoid of the parasites remains intact (C), rhoptries (R) and micronemes (M). Bar is 1 µm. (b) Detail of a spherical shape organism showing the separation of the inner membrane complex (IMC) from the plasmalemma disrupting the pellicular structure. R – Rhoptry. Bar is 1 µm. (**C**) Replication analysis of *gap45* KO parasites were performed 96 hours post induction. Parasites were allowed to invade for 1 h prior replication for 24 h. To quantify the replication rate the number of parasites per parasitophorous vacuole was determined. Mean values of three independent assays are shown ± SEM (**D**) Trail deposition assays with indicated parasites were performed by allowing the parasites to glide on FBS coated cover slips for 30 minutes prior staining with α-Sag1 antibody. The assays were performed 96 hours post induction ± 50 nM rapamycin. Analysis of the *gap45* knockdown (Frenal et al., 2010) was performed after induction with ATc for 96 hours as described previously (Frenal et al., 2010). Scale bars: 20 µm. (**E**) Quantification of trail-deposition assays on indicated parasites shows no effect on the number of gliding trails in *gap45* KO parasites. Mean values of three independent experiments are shown, ± SEM. The number of trails was standardised to wild type parasites. (**F**) Kinetics of gliding motility of *gap45* KO parasites. The average speed of 19 manually tracked parasites was analysed over a time interval of 30 minutes. (**G**) Indicated parasites treated with and without 50 nM rapamycin 96 h prior performing egress assays. Intracellular parasites were artificially induced with Ca^2+^ ionophore (A23187) to trigger egress for 5 min. For quantification mean values of three independent egress assays are shown ± SEM. The egress rates were standardised to wild type parasites. (**H**) Invasion assays were performed 96 h after excision of *gap45*. Parasites were allowed to invade for 1 hour prior washing the cover slip with 1 x PBS to remove all extracellular parasites. Parasites were fixed after 24 hours and the amount of vacuoles in 25 fields of view were determined. Mean values of three independent assays are shown ± SEM. All strains were standardised to wild type parasites. (**I**) Immunofluorescence analysis of invading parasites demonstrates that parasites lacking *gap45* are capable of invading the host cell forming a tight junction (TJ). The TJ was visualised by staining against RON4. Scale bar: 5 µm.

Quantitative analysis of parasite motility revealed that *gap45* KO parasites formed gliding trails similarly to control parasites (Figure 5D,E) and that unlike previously reported, *gap45* KD parasites were equally capable of gliding (Figure 5D,E). In addition, we analysed time lapse movies and found that depletion of GAP45 did not affect gliding speed as severely as seen in case of *myoA* KO since *gap45* KO parasites can move double as fast (0.5 µm/s) as *myoA* KO parasites (0.2 µm/s)(Figure 5F).

Interestingly, it seems that morphologically less affected parasites (note: MyoA and MLC1 are cytosolic independently of parasite morphology in *gap45* KO; Figure 5A) are capable to glide more efficiently, almost like control parasites. In contrast a slower, less efficient motility characterized spherical parasites (see also movies S6, 7).

Depletion of GAP45 resulted in a pronounced block of parasite egress (Figure 5G) similarly to what we observed for *mlc1* KO parasites but it caused a more severe block in host cell invasion than those seen for *myoA* KO or *mlc1* KO (∼7% when compared to control parasites, Figure 5H). Importantly, even morphologically affected, the spherical parasites were found to form a normal appearing TJ (Figure 5I), suggesting that key features of the invasion process were preserved.

Together these data demonstrate that GAP45 has a role in providing the IMC with its typical structure thereby promoting the anchorage of the MyoA motor complex in the IMC. However, since gliding motility was less affected, it appears that the parasite can efficiently produce forward movement in the absence of a myosin motor properly anchored along the IMC.

In addition, the significant loss of invasiveness is unlikely to result from an impairment of the gliding motility as previously suggested but is rather caused by the morphologic defect of these mutants.

While the study of the motor mutants demonstrates that alternative pathways must be in place that can drive gliding motility and invasion, it does not rule out a critical function of other myosin motors. However, in case of GAP45 this motor must be sharply localised at the apical tip of the parasite which is still intact, as demonstrated by ultrastructural analysis (Figure 5B). In order to investigate if invasion requires a myosin motor, we investigated the role of parasite actin in more detail, since any myosin will require polymerised actin in order to exert its function as a motor.

### Phenotypic analysis of act1KO parasites

We previously reported that parasite actin is essential for apicoplast replication and hence essential for the asexual life cycle of the parasite [12]. Despite that depletion of actin resulted in the loss of the apicoplast within 24 hours after removal of Act1, *act1* KO parasites continued to replicate although at a slightly slower rate (Figure 6A) before they died within vacuoles only ∼ 10 days after removal of the gene, [12,24].

**Figure 6.**
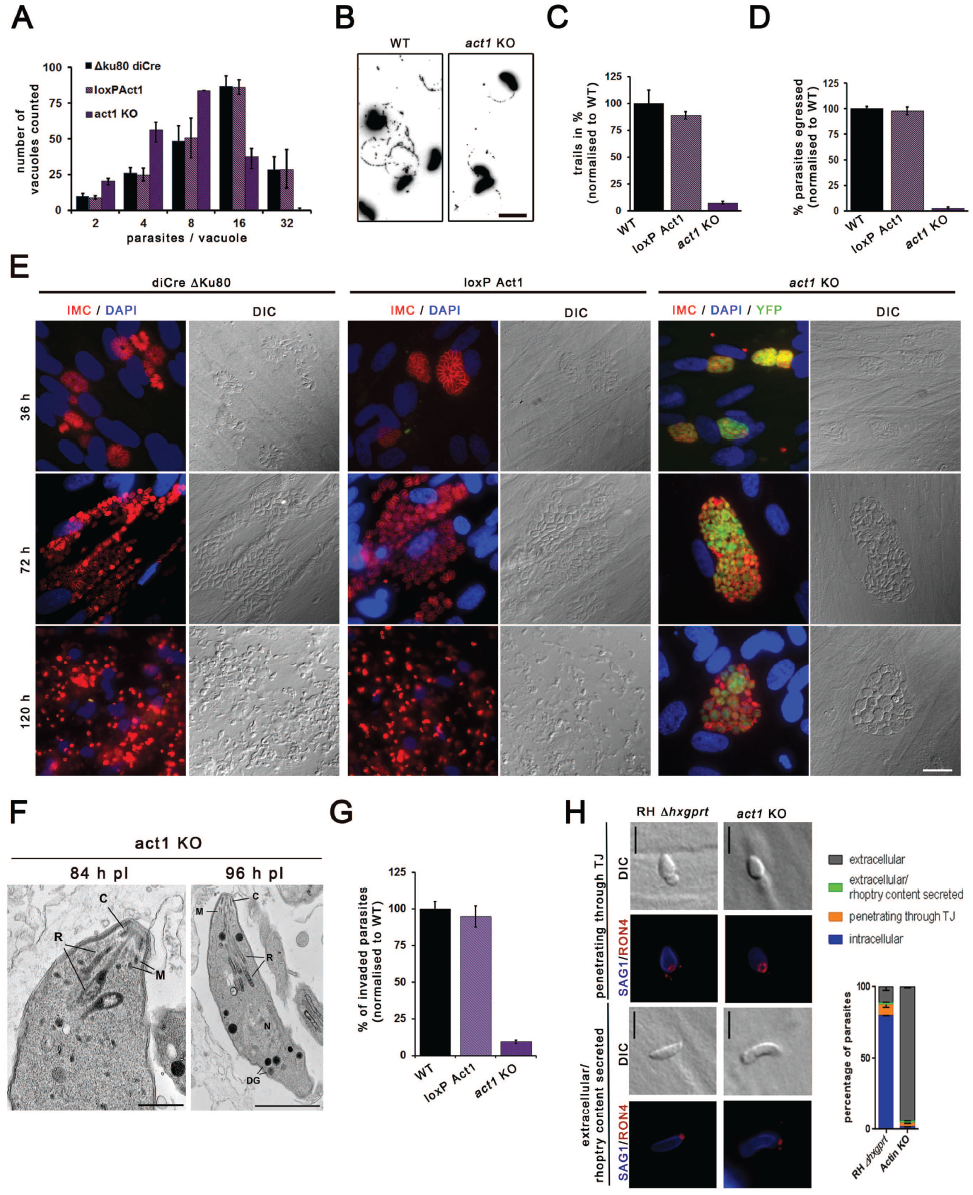
Generation and characterisation of *act1* KO (A) Replication analysis of act1 KO parasites was performed 96 hours post induction. Parasites were allowed to invade for 1 h prior intracellular growth for 24 h. To quantify the replication rate the number of parasites per parasitophorous vacuole was counted. Mean values of three independent assays are shown ± SEM **(B)** Trail deposition assays with indicated parasites were performed by allowing the parasites to glide on FBS coated cover slips prior staining with α-SAG1 antibody. Scale bars: 20 µm. **(C)** Quantification of trail-deposition assays on indicated parasites shows a significant reduction on the number of gliding trails in act1 KO parasites (10%). Mean values of three independent experiments are shown ± SEM. The number of trails was standardised to wild type parasites. **(D)** Indicated parasites were artificially induced with Ca2+ ionophore (A23187) to trigger egress for 5 min. For quantification mean values of three independent egress assays are shown ± SEM. The egress rates were standardised to wild type parasites. **(E)** Parasites were inoculated on HFF cells, fixed after 36 h, 72 h and 120 h and stained with α-IMC1 antibody. While the control parasite lines fully lysed the host cells after 120 hours, the act1 KO parasites were not capable of egressing the host cell and started dying within an intact parasitophorous vacuole. Scale bars represent 25 µm. **(F)** Ultrastructural appearance of *act1* KO parasites showing normal location and orientation of the apical organelles consisting of the conoid (C), rhoptries (R), micronemes (M) and dense granules (DG). N – nucleus. Bars represent 0.5 µm and 1 µm respectively. **(G)** Invasion assays of indicated parasites were performed 96 h after treatment with ± 50 nM rapamycin. 300 parasites were counted if they were invaded or still attached to host cells after 30 minutes of invasion time. Mean values of three independent assays are shown ± SEM. **(H)** IFA against the tight junction protein RON4 and SAG1 in act1 KO and RH Δhxgprt tachyzoites, scale bar represents 5 μm. Dot at apical tip of extracellular parasites visualises that rhoptry content has been secreted. Right graph depicts quantification of extracellular, extracellular and rhoptry content secreted, internalising and intracellular parasites. Data shows mean values ± SEM of three independent experiments in triplicate.

To analyse the involvement of actin during motility driven processes, parasites were incubated for 96 hours after activation of DiCre, which is 24 hours longer then the time required to reduce Act1 to undetectable levels as shown in immunoblot and immunofluorescence assays (Figure 1A, 1C). It is worth mentioning that identical phenotypes were observed at earlier time points (48 hours), when residual actin was still detected (not shown), indicating that once the actin concentration is below the critical concentration, the phenotype is absolute and cannot be enhanced further. Analysis of parasite motility in trail deposition assays revealed that approximately 10% of *act1* KO parasites were capable of forming trails (Figure 6B, C and Figure S2) that in most cases corresponded to half-circles or full circles, similar to *myoA* KO parasites. This residual motility was completely blocked, when parasites were incubated in high potassium (endo) buffer [18] (Figure S2). Despite maintaining a residual motility, the *act1* KO tachyzoites were blocked in host cell egress after Ca^2+-^Ionophore stimulation [25]; Figure 6D) and no spontaneous egress were observed over a 120 hours monitoring period (Figure 6E). Even though in some cases parasites within the parasitophorous vacuoles started to die at this late time point, artificial release of parasites from the host cell and subsequent inoculation on fresh host cells demonstrated that parasites can re-invade new host cells. Actin was recently suggested to be involved in positioning of rhoptries to the apical complex [26]. However, when the ultrastructure of artificially released *act1* KO parasites was analysed we found no morphological defects of the apical complex, in particular on rhoptry positioning, as previously reported [12]. Therefore, we conclude that the invasion defects cannot be attributed to morphological defects of the secretory organelles. It was found that the overall invasion rate was ∼10% of that control parasites (Figure 6G). Next the ability of parasites to form a TJ was analysed as described above for *myoA* KO parasites (Figure 2F). Although it has been reported that TJ formation is not blocked upon interference with parasite actin [13,27], we found that only 10% of all parasites were capable of establishing a TJ in pulse invasion assays, when compared to control parasites. This result fits well with the overall reduction in the invasion rate of *act1* KO parasites. It was thus concluded that *act1* KO are capable of penetrating host cells upon TJ formation (Figure 6H) and that a step upstream of TJ formation is blocked in absence of Act1.

## Discussion

Apicomplexan parasites have evolved unique organelles and structures to actively invade the host cell. The events leading to host cell invasion is a highly coordinated process that can be defined in several steps [8]: (i) parasite approach of the potential host cell by gliding (after host cell egress); (ii) host cell recognition and apical contact with the host PM; (iii); (iv) establishment of the tight junction (TJ) and finally (v) the entry into the host cell. It is widely accepted that the machinery that powers gliding motility and penetration into the host cell (v) is identical and depends on parasite actin, the MyoA motor complex and microneme proteins that act as force transmitters [28].

Conditional knock out is a powerful approach to decipher function(s) of a given protein by deleting gene in a tissue or time specific manner. First generation of these techniques has included the use of tetracycline-inducible transactivator system and has been successfully applied to *Toxoplasma*, in particular to the gliding machinery components [15,29–33]. However, in no case, the knockdown of one of these core components resulted in the expected block in host cell invasion and this discrepancy has been explained by leaky expression of the respective gene of interest.

The recent adaptation of a conditional recombination system in *T. gondii* has now allowed the complete removal of components of the assumed invasion machinery, such as *myoA, act1,* the micronemal proteins MIC2 and AMA1 [11,12] or the rhomboid proteases ROM4 (Abstract #175 Molecular Parasitology Meeting, Woods Hole 2013).

Intriguingly, all generated mutants for the core components of the invasion machinery (see summary Table 1) remained capable of invading the host cells despite the absence of elements thought to control the function of the actin/myoA-based motor and even in absence of actin itself. While compensatory or redundancy mechanisms are likely for some of these mutants, in particular for the myosins, these results also open the possibility of alternative molecular pathways to account for gliding motility and host cell invasion.

**Table 1.** **Overview of KO mutants and phenotypes for the gliding machinery**

| Parasite line | Viability in culture | Intracellular growth | Egress (%) standardised to WT | Invasion (%) standardised to WT | Gliding (%) standardised to WT | Organellar defects | Reference |
| --- | --- | --- | --- | --- | --- | --- | --- |
| **MyoA KO** | Viable | No effect | 2 % | 16 % | 37 % | No defect | Andenmatten et al., 2012 |
| **MyoB/C KO** | Viable | No effect | No effect | 100% | 100% | No defect | This study |
| **MyoA/B/C KO** | N.d. | No significant effect | 2 % | 5 % | N.d. | N.d. | This study |
| **MLC1 KO** | Up to 14 days | No effect | 5 % | 28 % | 42 % | No defect | This study |
| **GAP45 KO** | Up to 14 days | No effect | 4 % | 6 % | 100% | IMC looses contact to PM, MyoA, MLC1 cytosolic | This study |
| **GAP40 KO** | Not viable | Blocked | N.d. | N.d. | N.d. | N.d. | In preparation |
| **GAP50 KO** | Not viable | Blocked | N.d. | N.d. | N.d. | N.d. | In preparation |
| **Act1 KO** | Up to 10days | Slightly slower | 2 % | 10% | 10 % | Apicoplast replication | This study |

The phenotypic analysis of *myoA* KO parasites suggested that MyoA plays a critical, yet not essential role for gliding motility of the parasite. Furthermore, MyoA is involved in efficient host cell entry, since many parasites enter the host cell in a slow stop-and-go fashion. Strikingly, parasites are capable to enter at similar speed as control parasites, demonstrating that the necessary force for rapid entry can be generated in absence of MyoA. Interestingly, a triple KO for *myoA,B/C* resulted in a more severe phenotype compared to single *myoA* KO, in strong support for overlapping functions of parasite myosins. Such interchange ability and redundancy has recently been demonstrated for mouse nuclear myosin I (NM1) that can be complemented by Myo1c [34]. When the MyoA-associated and -regulatory partner MLC1 was depleted, we observed a mislocalisation of, MyoA. As expected, MLC1 loss caused a similar phenotype as observed for *myoA* KO parasites including a reduced gliding motility with mainly circular trails detectable and comparable invasiveness through a normally appearing TJ. In addition host cell egress was completely blocked and therefore maintenance of *mlc1* KO parasites was not possible. Of note, since MLC1 depletion triggered MyoA mislocalisation, functional redundancy in the repertoire of myosin light chains is unlikely. In good agreement, depletion of MTIP in *Plasmodium berghei* results in degradation of MyoA [23]. However, based on these results the presence of a different motor complex that can substitute for MyoA, such as MyoD-MLC2 [21] cannot be excluded.

GAP45 depletion had a major effect on the shape of extracellular parasites that lost their typical crescent shape and rounded up. This morphological change was accompanied by the redistribution of MLC1 and MyoA to the cytosol of the parasites. While these results confirm previous results by Frenal *et al.* 2010, it was surprising to find that even morphologically disrupted (round) *gap45* KO parasites were capable to glide, albeit slower than controls. Strikingly, we find that *gap45* KO parasites glided more efficiently than *myoA* KO parasites, confirming that the motility can be generated in absence of the known motor complex. Although it is possible that a different, unknown myosin motor is involved in this process, one has to consider that the IMC, the platform for a potential second motor is disrupted in *gap45* KO parasites. Finally, invasion by GAP45 depleted parasites was significantly reduced, probably as a consequence of the morphological defect but not of the loss of gliding motility. Importantly, as described for *mlc1* KO and *myoA* KO parasites, host cell entry proceeded through a normal TJ. Similar to *mlc1* KO parasites long term cultivation of *gap45* KO parasites was not possible most likely because of a block in host cell egress.

Intriguingly, depletion of parasite actin did not result in a complete block of motility, since short circular trails were readily detected in motility assays, suggesting that a residual motility is possible even in absence of parasite actin.

As is the case for motility, depletion of the core components of the known invasion machinery did not result in a block of host cell invasion. Instead it appears that the major limitation caused by depletion of this machinery lies in the delayed formation of the TJ. In case of *myoA* KO parasites and *act1* KO parasites TJ formation was severely delayed, explaining a reduction in overall invasion rate. However, once the TJ was formed parasites entered the host cell regardless of the integrity of the typical invasion machinery. As demonstrated for *myoA* KO parasites, the entry process was less efficient, with many parasites moving into the host cell in a stop-and-go fashion. However, since some parasites could enter host cells at the same speed as control parasites, it is possible that the parasite or the host cell can generate the force required for entry. Together these results suggest that gliding motility is critical in a step upstream of TJ formation, but as soon as a TJ is formed the parasite can enter, albeit less efficiently, in absence of their actomyosin system.

Based on previous studies that implicate an important role of host cell actin during invasion [35–37] we favour the hypothesis that once the TJ is formed host cell actin plays an important role during invasion of the analysed mutants.

In summary, this study leads to three hypotheses. 1. Components of the invasion machinery show multiple redundancies. 2. A compensatory invasion mechanism is in place that can substitute for the loss of a functional actin-myosin-system. 3. **Our current model that predicts a linear motor for the generation of force for motility and invasion needs to be revised**.

In line with the third possibility, we propose that a gelation-solation osmotic engine (Figure 7) could drive the parasite propulsion, similar to the model proposed in [38]. Specifically, a gel of actin-like filaments and acidic protein micromolecules secreted by micronemes at the apical end of the cell would be coated by both immobile heavy and mobile light cations in the cytoplasm, as was shown for actin gels in vitro [39]. The mobile cations generate osmotic pressure that is balanced by tension of the elastic gel. Partial disassembly of the gel caused by degradation of its macromolecules will cause weakening of the gel elastic modulus and gel swelling. This swelling will push the leading edge of the cell forward, providing there are adhesions of the gel to the substrate that are spatially biased to the rear (Figure 7). Subsequent complete disassembly of the older weaker gel and reassembly of the dense gel at the new leading edge completes the protrusion cycle, which can be step-like if the assembly-disassembly is cyclic or smooth if the gel assembles and disassembles continuously. The rear of the cell will be pulled forward by the membrane tension generated by the protrusive force [40] and the cytoplasmic flow from the rear caused by the gradient of the pressure in the cell – such gradient is made possible by the poroelastic nature of the cell cytoskeleton [41,42]. Simple estimates in the supplemental material demonstrate that such gelation-solation osmotic engine is physically feasible. The idea of a macroscopic osmotic engine coupled with the gel elasticity was theoretically proposed and experimentally proved a long time ago [43]. Ability of osmotic gradients to propel membrane vesicles at speeds comparable with those reported in this paper was demonstrated experimentally [44]. The proposed hypothesis is also supported by findings that rapid dynamics of adhesions and fast cycles of assembly and disassembly of actin filaments are necessary for the fast motility of the parasite [45–47]. The fact that high potassium buffer completely blocks gliding motility [18] lends additional support for the model. The high cytoplasmic pressure in the motile parasite is evident from the rounded shape of the GAP45 mutant in our study and from the bleb-like protrusions evident in plasmodium ookinete mutants [48].

**Figure 7.**
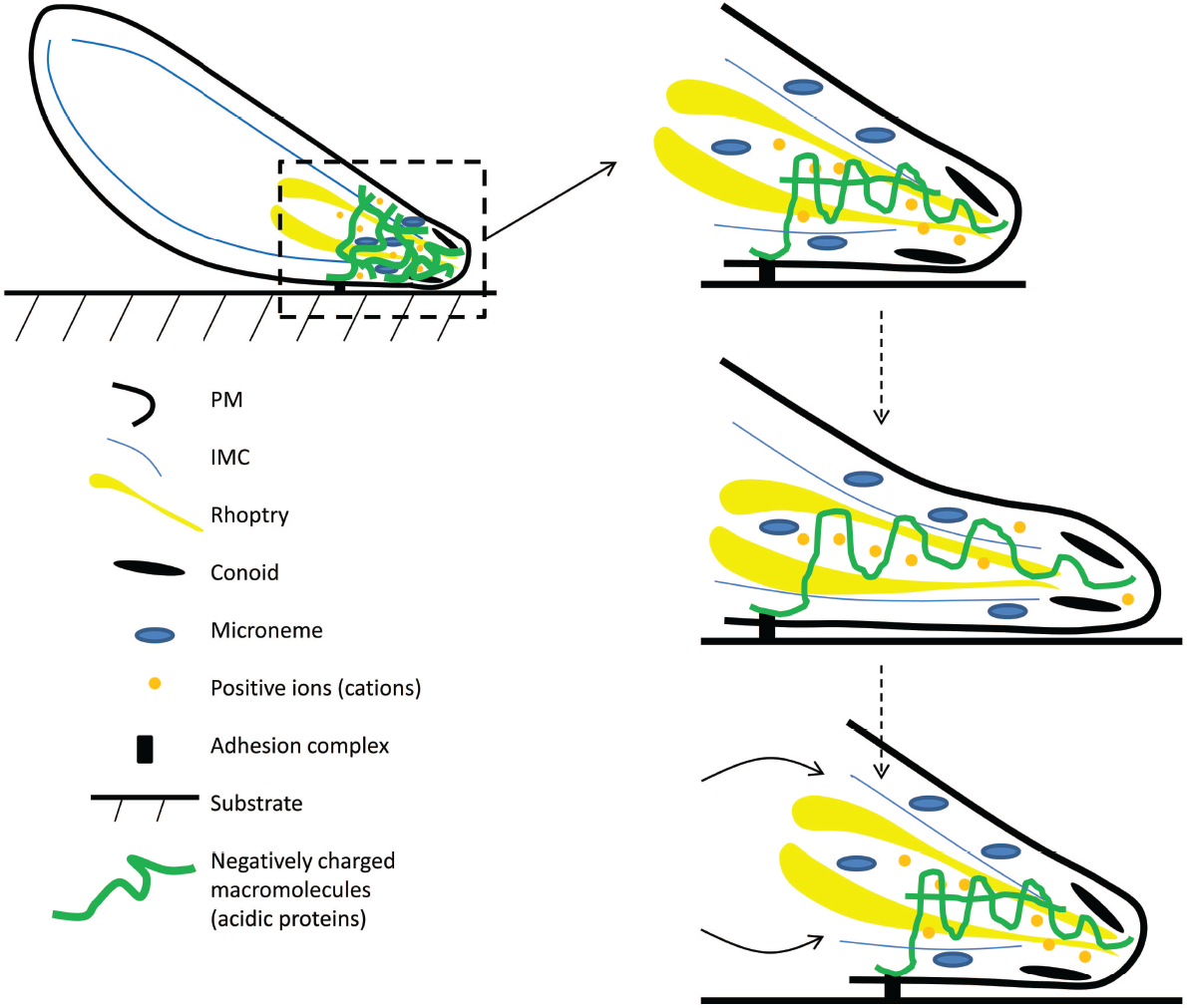
Gelation-solation model of the cell propulsion. Schematic side view of the cell. We hypothesize that there is a gel of negatively charged micromolecules, adhering to the substrate and surrounded by mobile cations, at the apical end of the cell. Initially, the gel is dense; osmotic pressure of the cations is resisted by effective elastic tension in the gel. The cycle of protrusion is illustrated at the right; for simplicity, only one long (coiled) and one short (almost straight) negatively charged macromolecules are shown (top). Middle: Partial gel disassembly (illustrated by disappearance of the short macromolecule) causes mechanical weakening of the gel. Osmotic pressure of the mobile cations, as a result, swells the gel (illustrated by the uncoiling of the long macromolecule). This gel swelling causes protrusion at the apical end (dashed) providing that adhesions are at the posterior end of the gel. Bottom: subsequent complete disassembly of the older macromolecules and adhesions and reassembly of the nascent dense gel and adhesions at the leading edge completes the cycle of protrusion. Membrane tension and volume conservation result in flow (arrows) and forward translocation of the whole cell.

In principle, other modes of parasite propulsion driven by the osmotic pressure are possible. First, it was shown theoretically that secretion of charged molecules at one side of the cell and their degradation at the other side coupled with water flow through permeable cell membrane can create water intake at the front and outflow at the rear and accompanying force that can propel the cell [49]. Sufficient membrane permeability probably requires aquaporins channels; however, aquaporin KO in *P. berghei* is viable and has no defect in gliding motility [50]. On the other hand, other transporters in the plasma membrane could be sufficient for the transmembrane water transport. Another possibility is that the inner membrane complex, transiently adhering to the substrate, acts as a piston separating the parasite’s tip and rear. Cycles of osmotic pressure increases and relaxations at the rear could then drive protrusion. Our observation that the *gap45* KO in which effective connection of the inner membrane complex to the plasma membrane and substrate is lost, but which is still motile, argues against this mechanism.

Lastly, as is clear from our data, MyoA plays an important role in accelerating the motility and making it more steady and persistent. It is tempting to speculate that actin filament polymerize directionally (perhaps the directionality is regulated by the ion gradient at the apical tip), and then MyoA molecules could be gliding and biased to the rear of the gel, where their interactions with adhesion molecules could ratchet the cycles of gel swelling into effective protrusions of the cell leading edge. Future experiments will test whether the hypothesized osmotic engine is indeed driving gliding motility.

## Materials and Methods

### Cloning of DNA constructs

All primers used in this study are listed in Supplementary Table 1. To generate *loxPMLC1loxP–YFP*–*HX* the *mlc1* 3′UTR was amplified from genomic DNA using the primer pair 3′UTR MLC1 fw/rv, and the PCR fragment was cloned into *p5RT70loxPKillerRedloxPYFP–HX* [12] via SacI. The *mlc1* ORF (TGME49_257680) was amplified from cDNA using the primers MLC1 ORF fw/rv, and was cloned into the parental vector using EcoRI and Pac1. To put *mlc1* under the control of the endogenous promoter a 2 kb fragment upstream of the start codon of *mlc1* was amplified from genomic DNA using the oligos 5′UTR MLC1 fw/rv and cloned into the parental vector using EcoRI and ApaI.

The *loxPGAP45loxP–YFP–HX* construct was generated using the strategy described for *loxPMLC1loxP–YFP–HX* with minor alterations. First, the *gap45* ORF (TGME49_223940) was amplified from cDNA using the primer pair GAP45 ORF fw/rv and digested with EcoRI/PacI. Next a 2 kb fragment upstream of the start codon was amplified using 5’UTRGAP45 fw/rv and digested with ApaI/EcoRI. Both ORF and endogenous were cloned at the same time point into the parental vector *p5RT70loxPKillerRedloxPYFP–HX*. Last the *gap45 3’UTR* was amplified from genomic DNA using the primer pair *3’UTR GAP45 fw/rv via PciI*.

To generate the *Myosin B/C KO–Bleo* construct, the bleomycin selection plasmid pBSSK+SAG1_BLE_SAG1 was used as parental vector [51]. First the *myob/c* 5’UTR was amplified from genomic DNA using the primer pair 5’UTR MyoB/C fw/rv and integrated into the parental vector via KpnI/HindIII. Afterwards the *myob/c* 3’UTR was cloned into the vector using the oligo pair 3’UTR MyoB/C fw/rv using PacI/SpeI.

### T. gondii cultivation, transfection and selection

*T. gondii* parasites (RH *Δhxgprt*) were grown in human foreskin fibroblast (HFF) and maintained in Dulbecco’s modified Eagle’s medium (DMEM) supplemented with 10% fetal calf serum, 2 mM glutamine and 25 µg/ml gentamycine. To generate stable transformants, 1 × 10^7^ of freshly lysed RH *Δhxgprt* –parasites were transfected with 60 μg DNA by electroporation. Selection based on mycophenolic acid (12.5 mg / mL in MeOH; MPA) / xanthine (20 mg / mL in 1M KOH; XAN) and phleomycin were performed as described earlier [52].

### Generation and verification of parasite lines

The conditional *mlc11* KO strain (*ku80::diCre/endogenous mlc1::loxPmlc1loxP*, referred to as loxPMLC1) was generated by transfecting 50 μg of the plasmid *loxPMLC1loxPYFP–HX* into the *ku80::diCre* [12] parasites to replace endogenous *mlc1*. After transfection parasites were selected for stable integration using XAN and MPA as described previously. Following the selection process the parasite pool was serially diluted to isolate single clones. The resulting loxPMLC1 strain carries only the Cre inducible copy of *mlc1*, allowing excision of *mlc1* upon rapamycin addition (50 nM in DMSO for 4 hours prior to washout) to generate the *mlc1* null mutant (*ku80::diCre/mlc1*^−^ referred to here as *mlc1* KO). Confirmation of 5′ UTR integration and site–specific recombination leading to the excision of *mlc1* was confirmed by PCR using the oligo set MLC1 5′UTR fw (1) and YFP rv (1’). The 3′ UTR integration into the correct locus was analysed using the primers HX fw2 (2) and MLC1 3UTR rv (2′).

To generate the conditional *act1* KO and *gap45* KO strains, 60 µg of the plasmids *loxPAct1loxPYFP-HX* and *loxPGAP45loxPYFP-HX* were transfected into a novel diCre Δku80 strain (Pieperhoff *et al.,* in preparation) respectively. Following transfection parasites were selected for stable integration using XAN and MPA as described previously [52]. After the selection process single clones were isolated. The resulting LoxPAct1 and LoxPGAP45 strains bear only the Cre inducible copy of *act11* or *gap45*, allowing excision of *act11* and *gap45* upon rapamycin addition (50 nM in DMSO for 4 hours prior to washout) to generate the *act1* and *gap45* KO lines (*diCre* Δ*ku80/act11*^−^, *diCre* Δ*ku80/gap45*^−^) referred to here as *act1* KO and *gap45* KO respectively). Confirmation of 5′ UTR integration and site–specific recombination of *Act1* was confirmed by PCR using the oligo set described previously [12]. Verification of 5′ UTR integration and excision of *gap45* was confirmed by PCR using the oligo pair GAP45 5′UTR fw (1) and YFP rv (1’). Correct integration into 3’ locus was analysed using the primers HX fw2 (2) and GAP45 3UTR rv (2′).

To generate the conditional triple KO for *myoA* and *myoB/C,* 60 µg of the plasmid *Myosin B/C KO–Bleo* were transfected into the loxPMyoA strain [12] and selected for stable integration using Phleomycin as described previously [51]. Single clones were isolated resulting in the strain LoxPMyoA MyoB/C KO in which the MyoB/C locus was altered in a way that *myob/c* was replaced by Bleomycin. Loss of the *myob/c* gene was confirmed by PCR using oligo set MyoB/C gene fw/rv (3-3’). To verify the correct integration into the 5’ and 3’ locus the oligo pairs 5’UTR MyoB/C fw/rv (1-1’) and 3’UTR MyoB/C fw/rv (2-2’) were used respectively.

### PCR, Western blotting and immunofluorescence analysis

For the extraction of genomic DNA from *T. gondii* to use as a PCR template, parasites were pelleted and afterwards washed in 1x PBS. DNA extraction was performed using Qiagen DNeasy blood and tissue kit according to manufacturer’s protocol.

Western blot samples were generated by pelleting extracellular parasites and incubating them with RIPA buffer (50 mM Tris–HCl pH 8; 150 mM NaCl; 1% Triton X–100; 0.5% sodium deoxycholate; 0.1% SDS; 1 mM EDTA) for 5 min on ice to lyse the parasites. Afterwards samples were centrifuged for 60 min at 14,000 rpm at 4 °C and laemmli buffer was added to the supernatant. Unless indicated otherwise 5×10^6^ parasites were loaded onto a SDS acrylamide gel and immunoblot was performed as described previously [53].

For immunofluorescence analysis, infected HFF monolayers grown on coverslips were fixed in 4% paraformaldehyde for 20 min at RT. Afterwards coverslips were permeabilised in 0.2 % Triton X–100 in 1x PBS for 20 min, followed by blocking (2% BSA & 0.2% Triton X–100 in PBS) for 20 min. The staining was performed using indicated combinations of primary antibodies for 1 h and followed by secondary Alexa Fluor 488, Alexa Fluor 350 or Alexa Fluor 594 conjugated antibodies (1:1000–1:3000, Invitrogen–Molecular Probes) for another 45 min, respectively.

### Equipment and settings

For image acquisition z–stacks of 2 μm increments were collected using a UPLSAPO 100 x oil (1.40NA) objective on a Deltavision Core microscope (Image Solutions – Applied Precision, GE) attached to a CoolSNAP HQ2 CCD camera. Deconvolution was performed using SoftWoRx Suite 2.0 (Applied Precision, GE). Image acquisition was also conducted using a 100x and 63x oil objective on a Zeiss Axioskope 2 MOT+ microscope attached to an Axiocam MRm CCD camera using Volocity software, Images were processed using ImageJ 1.34r software and Photoshop (Adobe Systems Inc., USA).

### Plaque assay

The plaque assay was performed as described before [15]. 200-500 freshly lysed parasites were inoculated on a confluent monolayer of HFF cells and incubated for 5 days. The HFF monolayer was washed in 1x PBS and fixed with 4% PFA for 20 min.

### Replication assay

Assay was performed as previously described [15]. In brief, 5×10^4^ freshly released parasites were allowed to invade for 1 hour. Subsequently, three washing steps for removal of extracellular parasites were performed. Cells were then further incubated for 24 h before fixation. Afterwards parasites were stained with α-IMC1 antibody and the number of parasites per vacuole was counted. Mean values of three independent experiments +/ SEM were determined.

### Invasion assay

In this study 2 different types of invasion assays were performed, invasion/replication or invasion/attachment. For the invasion/replication assay 5×10^4^ freshly released parasites were allowed to invade for 1 hour. Subsequently, three washing steps for removal of extracellular parasites were performed. Cells were then further incubated for 24 h before fixation. Afterwards parasites were stained with α-IMC1 antibody and the number vacuoles in 15 fields of view were counted. Mean values of three independent experiments +/ SEM were determined The invasion/replication assay was used to analysed induced and non-induced loxPGAP45 parasites. For the invasion/attachment assay 5×10^6^ freshly released parasites were allowed to invade for 30 min to 1 hour and then fixed with 4% PFA. Afterwards extracellular parasites were stained with α-SAG1 antibody and either the number of extra and intracellular parasites in 15 fields of view was scored (as in the case of *myoA* KO) or 300 parasites were analysed if they were attached or invaded into the host cell (this quantification method was used for all remaining conditional KOs). Mean values of three independent experiments +/ SEM were determined.

### Egress assay

5×10^4^ parasites were grown on HFF monolayers on coverslips for 24 h. Media was exchanged for pre–warmed, serum–free DMEM supplemented with 2 μM A23187 (in DMSO) in order to artificially induce egress. After 5 min cells were fixed and stained with α-SAG1 antibody. 200 vacuoles were counted for their ability to egress out of the host cells. Mean values of three independent assays +/ SEM were determined.

### Gliding assay

Trail deposition assays have been performed as described before [54]. Briefly, freshly released parasites were allowed to glide on FBS-coated glass slides for 30 min before they were fixed with 4% PFA and stained with SAG1.

### Time lapse microscopy

Time-lapse video microscopy was conducted with the DeltaVision^®^ Core microscope using a 40X objective lens for invasion analysis and 20X objective for gliding analysis. Normal growth conditions were maintained throughout the experiment (37 °C; 5% CO_2_). Images were recorded at one frame per second using SoftWoRx^®^ software. Further image processing was performed using ImageJ 1.34r software.

In order to assess penetration time, heavily infected HFFs were scratched and the cells suspension was passed through 26 gauge needle three times to release parasites artificially. About 1 × 10^6^ parasites were then added onto HFFs grown in Ibidi µ-Dish^35mm, high^. Invasion events were observed after about 20 min when parasites had settled and corresponding penetration time was determined. To assess the gliding kinetics parasites were prepared akin to the trail deposition assay. Briefly, extracellular parasites were pelleted, washed in pre-warmed PBS and resuspended to a concentration of 1 × 10^6^ per 800 µl in pre-warmed HBSS. Parasites were then added to FCS-coated glass dishes (Ibidi µ-Dish^35mm, high^) and gliding was recorded for 30 min after parasites have settled. For analysis, 20 parasites were manually tracked using the manual tracking plugin for ImageJ 1.34r software and total displacement as well as average speed were determined.

### Staining and quantification of tight junction

Rhoptry secretion and tight junction formation was assessed as described by [55] with minor changes Briefly, 36 hours post inoculation heavily infected HFFs were scratched and parasites were released by passing the cell suspension through a 26 gauge needle followed by centrifugation at 600 g for 5 min. Supernatant was removed and parasites were resuspended in 3 × 10^6^/200 µl Endo buffer (44.7 mM K_2_SO_4_, 10 mM Mg_2_SO_4_, 106 mM sucrose, 5 mM glucose, 20 mM Tris, 0.3 5% (w/v) BSA, pH 8.2). Parasites were incubated at RT for 10 min before adding 200 µl to a confluent monolayer of HFFs grown on coverslips. Parasites and cells were incubated for an additional 3 min at RT followed by 20 min at 37 °C and 5% CO_2_ to let the parasite settle naturally. Please note that parasites are unable to secrete their micronemes in high potassium buffer, thus attachment is only lose. Therefore, supernatant was carefully replaced by 500 µl pre-warmed growth media and parasites were let to invade for 5 min at normal growth conditions. After three washing steps with PBS, the cells were then fixed with 4% paraformaldehyde followed by immunostaining under non-permeabilising conditions with rabbit α-Toxoplasma and mouse α-RON4 primary antibody followed by Alexa Fluor secondary antibody, respectively. Alexa Fluor 488/594 secondary antibody was used at 1:500 dilution to increase the RON4 signal at the tight junction. Assay was performed in triplicate at three independent occasions and 200 parasites were counted and calculated as percentage.

### Electron Microscopy

Samples of extra- and intra-cellular tachyzoites of wild type and GAP45 KO cultured for 48 and 72 hours were fixed with 2.5% gluteraldehyde in 0.1 M phosphate buffer pH 7.4 (1M Na2HPO4, 1M NaH2PO4). Samples were then processed for routine electron microscopy as described previously [56] In summary, samples were post-fixed in osmium tetroxide, en bloc stained with unranyl acetate, dehydrated, treated with propylene oxide and embedded in Spurr’s epoxy resin. Thin sections were stained with uranyl acetate and lead citrate prior to examination in a JEOL 1200EX electron microscope.

## Acknowledgement

We thank Gary Ward (University of Vermont), C.J. Beckers (University of North Carolina, Chapel Hill), A. Scherf (Pasteur Institute, Paris), J.F. Dubremetz (University of Montpellier), Dominique Soldati (University of Geneva) and John Boothroyd (University of Stanford) for sharing reagents. Gary Ward and Friedrich Frischknecht for fruitful discussions. This work was supported by the Wellcome Trust. M.M. is funded by a Wellcome Trust Senior Fellowship (087582/Z/08/Z) and an ERC Starting Grant (ERC-2012-StG 309255). N.A. is supported by an EviMalaR (European FP7/2007-2013, grant number 242095) PhD fellowship. The Wellcome Trust Centre for Molecular Parasitology is supported by core funding from the Wellcome Trust (085349).

## Supporting Information

**Estimates of the physical feasibility of the gelation-solation osmotic engine.**

Simple estimates demonstrate that the hypothesized gelation-solation osmotic engine is physically feasible. The protrusion force can be roughly estimated with Van’t Hoff formula [1]: osmotic pressure at the leading edge P ∼ cRT, where c is the molar concentration of the mobile cations, R is the gas constant and T is the absolute temperature. Assuming that c is on the order of ten micromolar (characteristic concentration of actin monomers, oligomers, short filaments and their counterions), this formula gives P ∼ 10–100 pN/mμ^2^, which is lower but comparable with the protrusive force at leading edge of migrating cells [2]. Gel elastic modulus originating from ion-mediated electrostatic interactions and entanglement of semi-stiff polymers [3,4] can be on the order of 100 pN/mμ^2^ and thus withstand such pressure; upon partial disassembly such gel relaxes [5] and deforms (by up to microns) enough to cause the gel swelling. The Darcy permeability of the gel to the cytoplasmic flow [6], K ∼ 0.01 mμ^4^/(pN×s) is high enough to allow the flow rate V ∼ K×P/L ∼ 1 mμ/s (L ∼ 1 mμ is the characteristic length scale at the leading edge) that will not limit the pathogen propulsion.

---

**Supplementary Figure 1:**
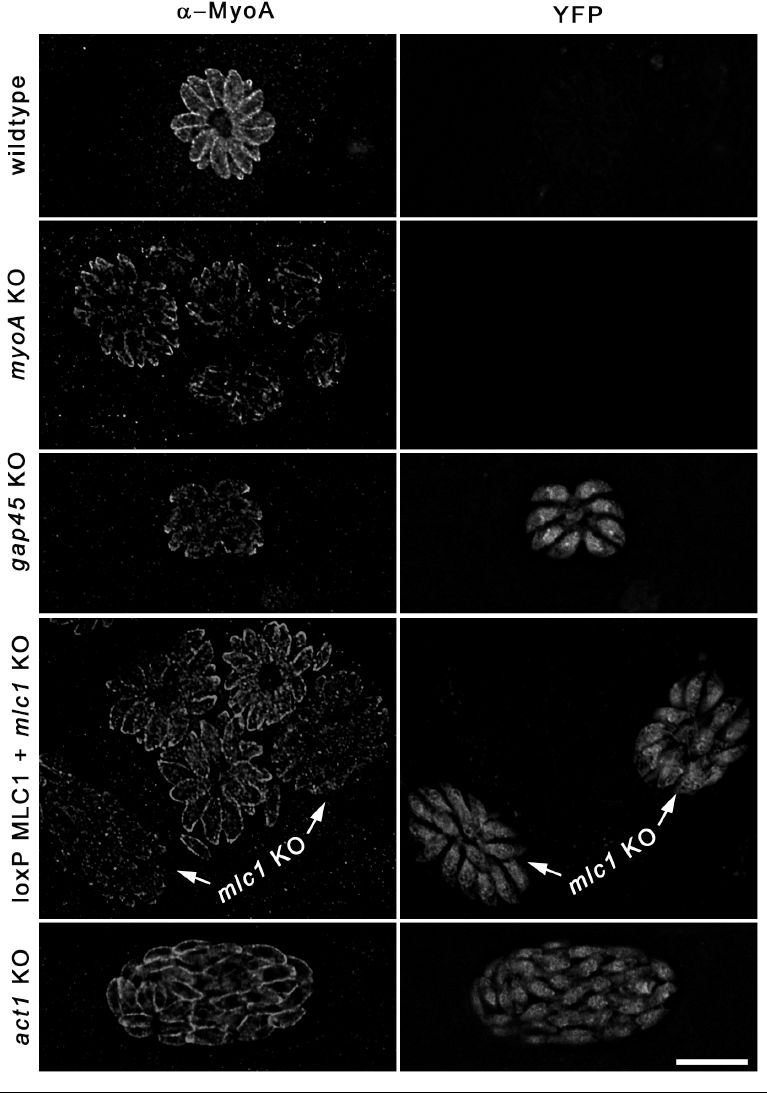
Comparison of MyoA antibody in various strains. Using the MyoA antibody for different strains revealed an unspecific background level within *myoA* KO parasites. Compared to wildtype parasites the signal was not detected at the periphery of the parasites and seemed mainly concentrated at the apical region of the parasites. Although no conclusion can be drawn concerning a possible depletion of MyoA in *mlc1* KO parasites, a clear localisation change (periphery to cytosol) of MyoA within *mlc1* KO and *gap45* KO can be observed. Scale bar: 10 µm.

**Supplementary Figure 2:**
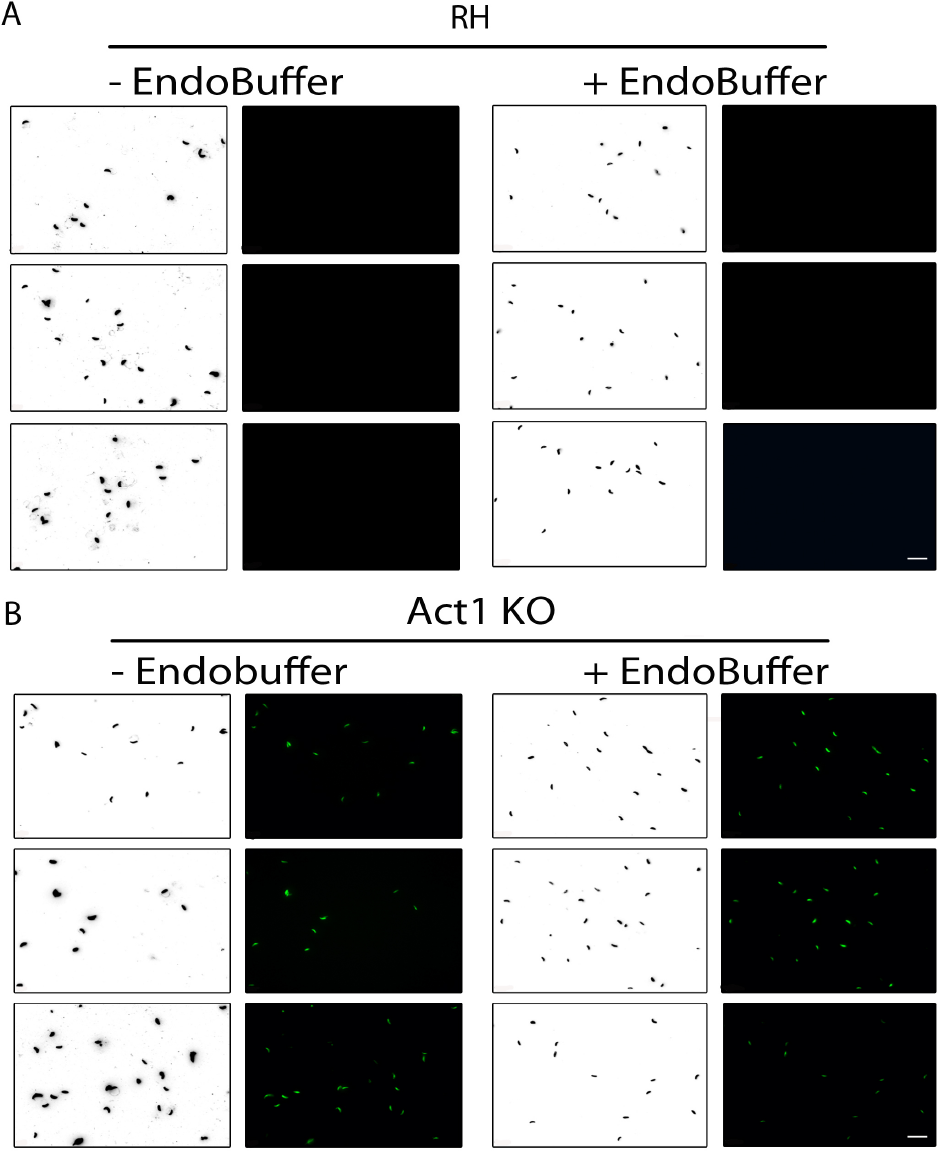
Gliding motility in absence of Act1. Trail deposition assay of RH Δ*hxgprt* (A) and *act1* KO (B) parasites. Parasites were allowed to glide either in gliding or endo buffer (high potassium buffer). *Act1* KO parasites were analysed 96 hours post rapamycin induction (as indicative by green fluorescence). In both cases no gliding trails were observed in Endo Buffer. Scale bar: 20 μm.

**Supplementary Movie 1: Circular gliding of RH *Δhxgprt***. Gliding on FCS coated glass bottom dishes was recorded at one frame per second and played at 7 frames per second, numbers indicate time in minutes:seconds and scale bars represent 10 μm.

**Supplementary Movie 2: Circular gliding of *myoA* KO**. Gliding on FCS coated glass bottom dishes was recorded at one frame per second and played at 7 frames per second, numbers indicate time in minutes:seconds and scale bars represent 10 μm.

**Supplementary Movie 3: Penetration of RH *Δhxgprt*.** Live time lapse microscopy of RH *Δhxgprt* invading an HFF cell. Frames were recorded every second and played at 5 frames per second, numbers indicate time in minutes:seconds and scale bars represent 5 μm.

**Supplementary Movie 4: Penetration of *myoA* KO with comparable kinetics to RH *Δhxgpr*t.** Live time lapse microscopy of *myoA* KO invading an HFF cell. Frames were recorded every second and played at 5 frames per second, numbers indicate time in minutes:seconds and scale bars represent 5 μm.

**Supplementary Movie 5: Penetration of *myoA* KO with slow kinetics** Live time lapse microscopy of *myoA* KO invading an HFF cell. Frames were recorded every second and played at 5 frames per second, numbers indicate time in minutes:seconds and scale bars represent 5 μm.

**Supplementary Movie 6: Circular gliding of crescent shaped *gap45* KO.** Gliding on FCS coated glass bottom dishes was recorded at one frame per second and played at 7 frames per second.

**Supplementary Movie 7: Circular gliding of spherical shaped *gap45* KO.** Gliding on FCS coated glass bottom dishes was recorded at one frame per second and played at 7 frames per second.

**Table S1.**
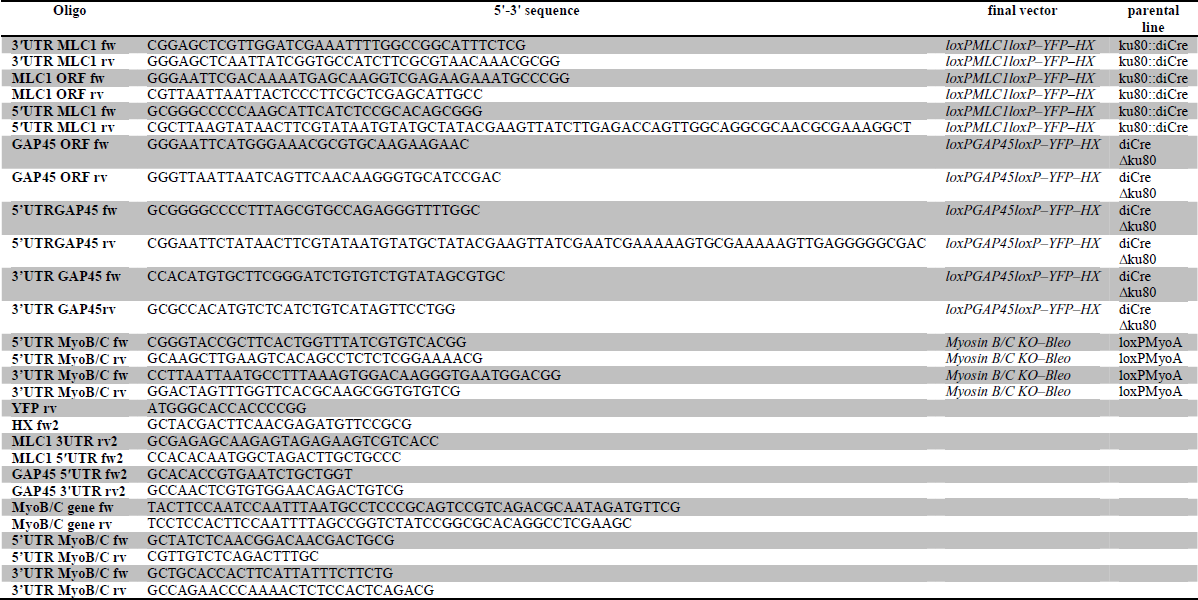
Summary of oligonucleotides used in this study.

